# Exploration of Bacteriophages Against Vancomycin-Resistant *Enterococcus faecium*: A Report from India

**DOI:** 10.64898/2026.08.03.742380

**Authors:** Lav Kumar Jaiswal, Nisha Rathor, Mrunalini Sonne, Minakshi Sahu, Chitra Nehra, Mahaveer Singh, Tanu Sagar, Rama Chaudhry

## Abstract

The vancomycin-resistant *Enterococcus faecium* (VREfm) has been declared as a high priority pathogen by World Health Organisation (WHO). It presents a major therapeutic challenge to healthcare, remaining with limited antibiotic options. Bacteriophage therapy has emerged as a promising alternative for combating antimicrobial-resistant pathogens. This study reports the isolation and characterization of bacteriophages active against VREfm and MDR *E. faecium* clinical isolates from India. Two bacteriophages, NIMS_EF375_N69_P9 (Φ1) and NIMS_EF375_N74_P12 (Φ2), were isolated from sewage by enrichment using MDR *E. faecium* strain EF375 as the propagation host. Lytic activity was confirmed by spot assay and double-layer soft agar plaque assay; both phages produced clear plaques of 1.5-2.0 mm. Transmission electron microscopy showed icosahedral heads with long non-contractile tails, presenting siphovirus-like morphotype within the class *Caudoviricetes*. Host range was assessed against 10 MDR *E. faecium* isolates (including the propagation host), 2 of which were VREfm. Φ1 lysed 5 of 10 isolates, including the VanA-type VREfm, whereas Φ2 lysed 7 of 10, including both VanA- and VanB-type VREfm. In time-kill assays against EF375 at an MOI of 1, Φ1 produced effective decline in optical density sustained through 24 h (∼80% reduction relative to the untreated control), whereas Φ2 suppressed bacterial growth with only ∼50% reduction. To the best of our knowledge, this study represents the first report from India on the isolation and characterization of bacteriophages active against clinical VREfm isolates. The broader host range of Φ2 and the stronger killing kinetics of Φ1 suggest complementary roles in a phage cocktail, warranting genomic characterization and *in vivo* evaluation.

## Introduction

AMR (Antimicrobial Resistance) is one of the most significant global challenges of the 21^st^ century. In particularly vancomycin-resistant *Enterococcus faecium* (VREfm) is a major cause of healthcare-associated infections globally. In 2017, VREfm was listed by the World Health Organization (WHO) as a high-priority pathogen, with limited treatment options and increasing global prevalence [1,2]. AMR is a natural and inevitable evolutionary phenomenon; it was first recognized shortly after the widespread introduction and mass production of penicillin [3]. The rapid development of antibiotics during the 1950s and 1960s initially helped to counteract the emergence of resistance however, the extensive and often indiscriminate use of antimicrobial agents, coupled with a marked decline in the discovery of novel antibiotics, has contributed significantly to the current AMR crisis [4,5]. Historically, *Enterococcus faecalis* was the predominant etiological agent of enterococcal infections. Since the 1980s, however, the incidence of *Enterococcus faecium* (*E. faecium)* infections has increased markedly and now accounts for approximately 50% of all enterococcal infections worldwide [6-10]. This shift has resulted in the widespread emergence of vancomycin-resistant VREfm strains [11]. Globally,*E. faecium* infections were estimated to cause approximately 11,000 deaths globally in 2021, with the burden projected to continue rising through 2050 [12].

In India, the first VREfm clinical isolate was reported by Mathur et al. in 1999 [13]. Since then, the prevalence of vancomycin-resistant enterococci (VRE) has increased substantially over the past two decades, with an overall pooled prevalence of 12.4%. Importantly, vancomycin-resistant *Enterococcus faecium* (VREfm) represents 58.2% of all VRE isolates, emphasizing its growing clinical burden in the country [2]. The vancomycin resistant *Enterococcus* species has been classified as high priority pathogen as per the Indian Priority Pathogen List (IPPL) [14], requiring urgent attention for the development of alternative therapeutic strategies. *E. faecium* is a Gram-positive, catalase-negative, non-spore-forming, facultative anaerobic bacterium, commonly present in diverse niches, including the gastrointestinal tracts of humans and animals, as well as environmental reservoirs such as water, soil and plants [2]. Although enterococci are normal commensals of the intestinal microbiota, they behave as opportunistic pathogens capable of causing urinary tract infections, bacteremia, infective endocarditis, meningitis, sepsis, surgical site infections, and device-associated infections, particularly among hospitalized and immunocompromised patients [15-21]. The clinical importance of *E. faecium* is particularly evident in intensive care units, where infections are associated with high morbidity and mortality [19,22].

The increasing prevalence of VREfm is largely attributed to the acquisition of transferable vancomycin resistance determinants. To date, nine vancomycin-resistance gene clusters (vanA, vanB, vanC, vanD, vanE, vanG, vanL, vanM, and vanN) have been identified in *Enterococcus* spp., with vanA and vanB being the most prevalent and clinically significant genotypes [23]. *E. faecium* possesses intrinsic resistance to several antimicrobial classes, including cephalosporins, low-level aminoglycosides, lincosamides, and trimethoprim-sulfamethoxazole, and readily acquires additional resistance to glycopeptides, β-lactams, aminoglycosides, oxazolidinones, daptomycin, and other clinically important antibiotics [24,25]. Consequently, VRE infections are associated with increased mortality, prolonged hospitalization, and greater healthcare costs. Furthermore, the ability of *E. faecium* to form biofilms further complicates treatment and contributes to persistent infections [26-28].

Several alternatives to conventional antibiotics have been explored to combat antimicrobial resistance, including predatory bacteria, bacteriocin, and bacteriophage therapy. Among these, bacteriophage therapy has emerged as a promising therapeutic approach and has gained renewed interest globally [29]. In contrast to antibiotics (broad spectrum), bacteriophages are host specific, targeting only particular bacterial species or strains. This specificity minimizes disruption of the commensal microbiota and reduces the selective pressure that drives the emergence and dissemination of antimicrobial resistance [30,31]. A large number of studies have demonstrated the efficacy of bacteriophages against a range of MDR bacterial pathogens for patient care as well as surface disinfection [32,33]. However, relatively limited scientific literature is available to explore and develop phage-based therapeutic strategies against. *E. faecium*.

At present (July 2026), a total of 134 articles (108 research and 26 review articles) related to phage targeting *E. faecium* are indexed in the NCBI PubMed database, of which only 10 publications specifically report bacteriophages targeting VREfm. India has contributed only five original research articles and six review articles on *E. faecium* bacteriophages, while no study from India has reported bacteriophages targeting VREfm. To the best of our knowledge, this is the first study from India describing the isolation and characterization of bacteriophages active against vancomycin-resistant *Enterococcus faecium* (VREfm). Accordingly, the present study reports the isolation and characterization of two lytic *E. faecium* bacteriophages and evaluates their biological characteristics and therapeutic potential against vancomycin-resistant and MDR and clinical isolates of *E. faecium*.

## Materials and Methods

### Host strain and culture conditions

The bacterial clinical isolates used in the present study were grown from various clinical samples, including urine, blood and pus. Isolates were identified as *E. faecium*, their antibiotic susceptibility patterns were determined using the VITEK-2 system, with MICs interpreted according to CLSI breakpoints [34]. The high level of vancomycin resistance was confirmed by Ezy MIC™ strip for vancomycin (0.016–256 μg/mL) (HiMedia Laboratories Pvt. Ltd). Clinical isolates of *E. faecium* were grown in Brain Heart Infusion (BHI) broth or BHI agar for bacteriophage isolation and characterisation. Enterococcal isolates showing resistance to both vancomycin and teicoplanin were classified as the VanA phenotype, whereas isolates resistant to vancomycin but susceptible to teicoplanin were classified as the VanB phenotype [35]. *Enterococcus faecium* isolates demonstrating resistance to three or more classes of antimicrobial agents were considered multidrug-resistant (MDR) [36].

### Isolation and propagation of bacteriophage

The bacteriophages infecting *E. faecium* were isolated from untreated wastewater collected from a sewage system. The isolation and purification of phages were performed as described earlier, with slight modifications [37-39]. Briefly, 10 mL of sewage was suspended in 40 mL of SM buffer (HiMedia Laboratories Pvt. Ltd). A 0.5 mL of chloroform (HiMedia Laboratories Pvt. Ltd) was added to the suspension, then spun down at 10,000 rpm for 10 min at 4°C. The supernatant was collected and filtered using 0.22 μm membrane filter. For phage enrichment, 10 mL of the filtered sample was pooled with an equal volume of double-strength Brain Heart Infusion (BHI) broth (HiMedia Laboratories Pvt. Ltd) and inoculated with 1 mL of an overnight culture containing *E. faecium*, incubated overnight at 37°C with continuous shaking. The next day, chloroform was added, and the supernatant was collected after centrifugation. The above supernatant was serially diluted and antibacterial activity was visualized by spot test followed by plaque assay using the double-layer soft agar method [40].

### Transmission Electron Microscopy (TEM)

TEM was performed for the morphological characterization of the bacteriophages. The phage lysates (10^8^-10^9^ PFU/mL), were filtered through 0.22 μm pore-size membrane filters. The filtrate was subsequently centrifuged at 25,000 × g for 75 min and the resulting pellet was washed thrice with 0.1 M ammonium acetate solution (pH 7.0). After removal of the supernatant, the pellet was again resuspended in ammonium acetate solution. Subsequently, 10 μL of the purified phage suspension was applied onto carbon-coated copper grids and allowed to adsorb for 10 min, stained by 1% uranyl acetate for 15 minutes and examined using transmission electron microscopy at the Sophisticated Analytical Instrumentation Facility (SAIF), AIIMS, New Delhi, India [41]. The size of both the bacteriophages was measured by ImageJ software.

### Host range determination

The phages were originally isolated against MDR *E. faecium* strain EF375 and tested on 2 VREfm and 7 other MDR *E. faecium*. A bacterial suspension with an OD_600_ of ∼0.5-0.6 was spread using the double-layer soft BHI agar method. After 10 µL of phage suspension (10^8^ PFU/mL) was spotted onto the bacterial lawn, the plates were then incubated for overnight at 37°C [42]. Lytic activity was then observed based on clear zones on the bacterial lawn.

### Phage Time-Kill Assay

For evaluating the ability of a bacteriophage to kill bacteria in a liquid culture Time-Kill Assay was performed which monitors the reduction of bacterial density over time. In the phage kill assay, *E. faecium* (0.2 -0.3 OD) was mixed with an equal amount of phage at an MOI of 1 and bacterial growth was quantified at 600 nm [43]. A 96-well microtiter plate was used for the experiment. BHI media without bacteria used as a negative control and inoculated bacteria without phage served as positive control. The plate was incubated at 37 ºC and the OD_600_ was monitored every 30 min. for 24 hrs in SPECTROstar^Nano^(BMGLABTECH, Germany). The measured experiments were performed in triplicates.

## Results

### Isolation and plaque morphology of the novel *Enterococcus faecium* bacteriophage

The Enterococcus viruses NIMS_EF375_N69_P9 and NIMS_EF375_N74_P12 were isolated using multidrug-resistant *E. faecium* strain EF375 as the host bacterium, which was cultured from urine sample. The phages NIMS_EF375_N69_P9 and NIMS_EF375_N74_P12 abbreviated as Φ1 and Φ2, respectively, for this paper. Plaque morphology was evaluated using the double-layer agar (soft agar overlay) assay. Both phages produced clear and distinct plaques on the lawn of *E. faecium* EF375 following incubation at 37 °C for 24 h. The average plaque diameter of both bacteriophages ranged from 1.5 to 2.0 mm **(Fig. 1. (a) and (b)**, indicating their lytic nature.

**Fig. 1.**
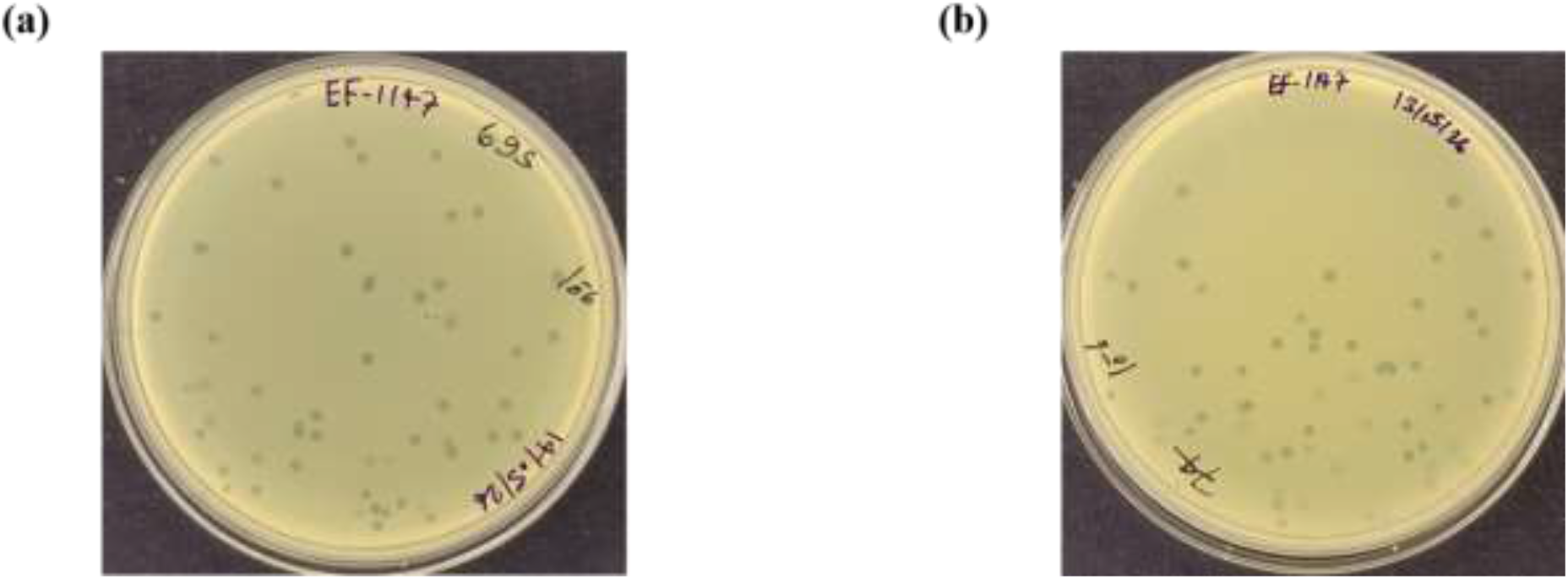
Plaques morphology of Enterococcus virus formed on the parental multidrug-resistant *E. faecium* strain EF375 lawn following the double-layer agar (DLA) assay and overnight incubation at 37 °C. **(a)** Clear plaques of 1.5-2.0 mm diameter produced by Enterococcus virus Φ1 **(b)** Clear plaques of 1.5-2.0 mm dimeter produced by Enterococcus virus Φ2.

### Transmission Electron Microscopy (TEM)

Transmission electron microscopy (TEM) analysis revealed that both phages possessed an icosahedral head and a non-contractile tail. *Enterococcus* virus Φ1 showed an icosahedral head (diameter: 50.68 ± 0.76 nm) connected to a long, non-contractile tail of 311.49 ± 2.61 nm (Fig. 2a). The *Enterococcus* virus Φ2 revealed an icosahedral head with a head diameter of 55.73 ± 0.05 nm, and a tail of 208.84 ± 1.73 nm in length (Fig. 2. b) (based on the 200 nm scale bar). The observed morphology is characteristic of *Siphovirus*-like bacteriophages belonging to the class *Caudoviricetes*.

**Fig. 2.**
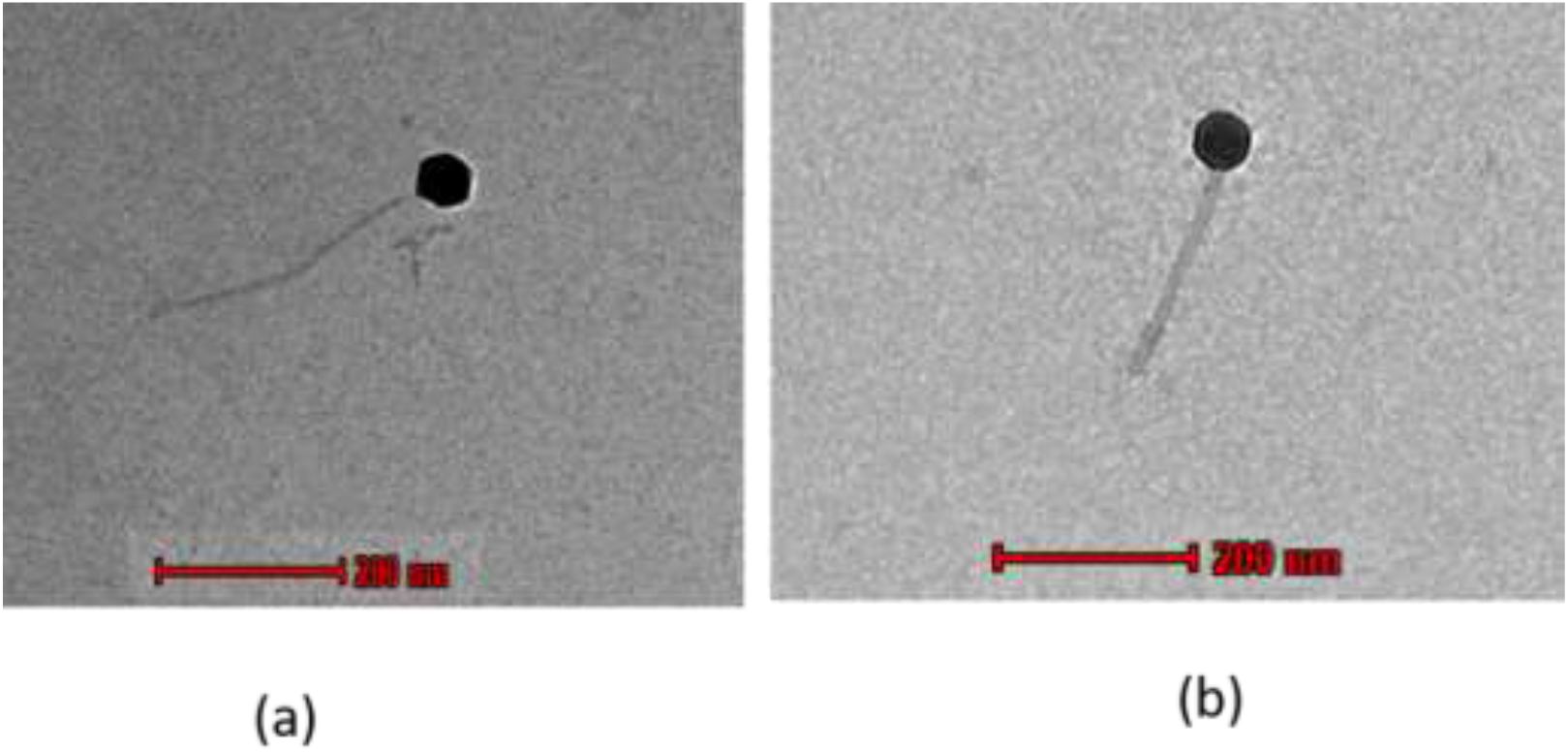
**(a)** Transmission Electron Microscopy-based negatively stained morphological analysis revealed that the Enterococcus virus Φ1 possess an icosahedral head with a diameter of 50.68 ± 0.76 nm and a tail of 311.49 ± 2.61 nm and **(b)** The phage Enterococcus virus Φ2 possess an icosahedral head with a diameter of 55.73 ± 0.05 nm, while the tail length was estimated to be 208.84 ± 1.73 nm based on the 200 nm scale bar.

### Bacterial Clinical isolates and Host range determination

The bacteriophages were initially isolated against MDR *E. faecium* (375), and subsequently tested on 2 VRE and 7 MDR *E. faecium* strains (Table 1). The host range analysis demonstrated that both isolated bacteriophages were capable of lysing *E. faecium* isolates with diverse antimicrobial resistance profiles. Out of the 10 *E. faecium* isolates tested (including parental host), Enterococcus virus Φ1 effectively lysed 1 type A VREfm and 4 MDR *E. faecium* isolates, whereas Enterococcus virus Φ2 lysed 2 VREfm (one VanA and one VanB Phenotype) and *5* MDR isolates, representing a broad host range **(Fig. 4)** and **Table 2**. The results indicate that Enterococcus virus Φ2 exhibited broader infectivity against the tested *E. faecium* collection compared to Enterococcus virus Φ1.

**Table 1.**
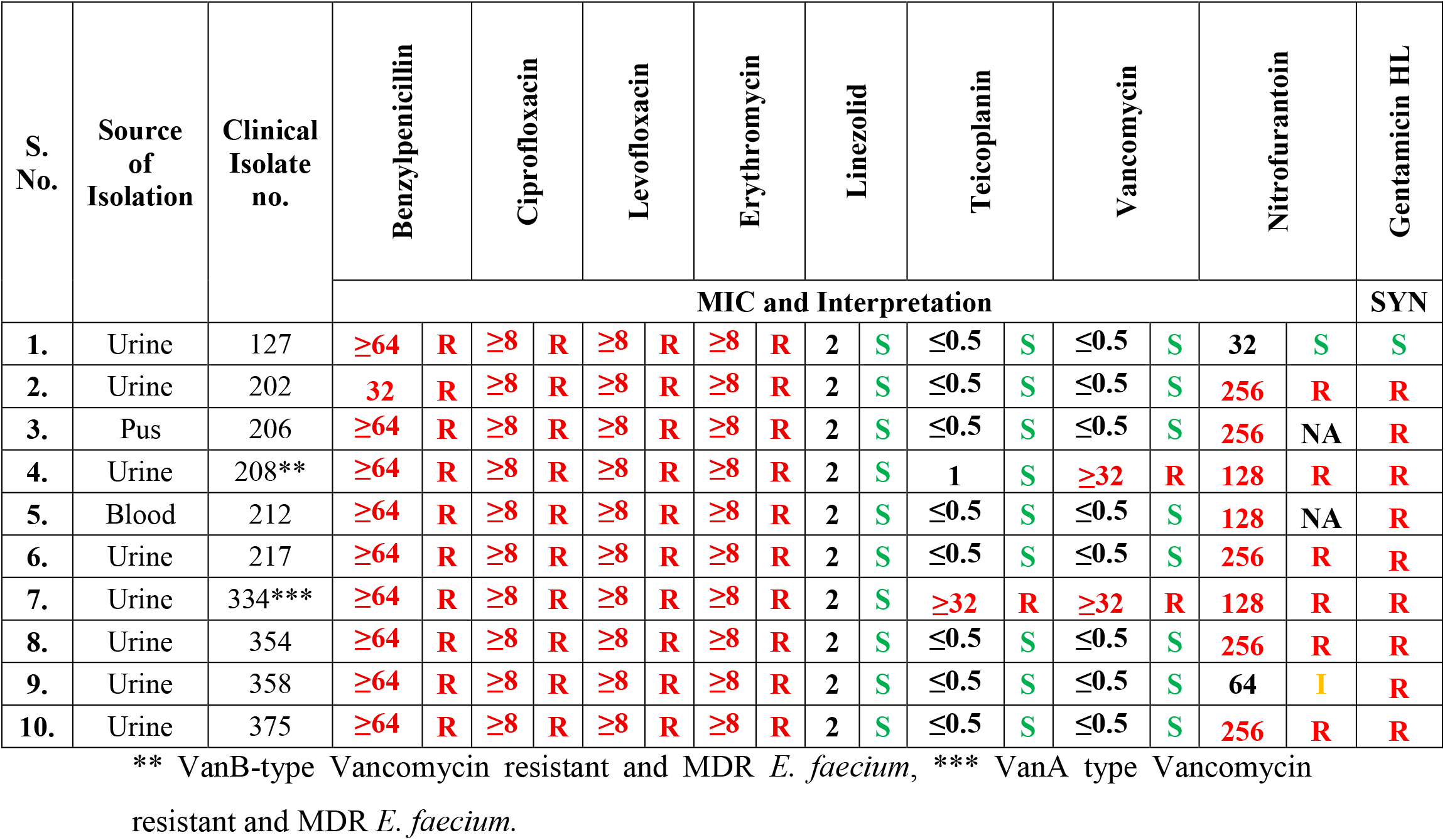
Antibiotic susceptibility profiles of MDR clinical Isolates of *E. faecium*. The MIC breakpoint of Enterococcus spp. according to CLSI 2025 [34] as for Penicillin MIC ≥ 16µg/mL, ciprofloxacin MIC ≥ 4µg/mL, levofloxacin MIC ≥ 8µg/mL, erythromycin MIC ≥ 8µg/mL, linezolid MIC ≥ 8µg/mL, teicoplanin MIC ≥ 32µg/mL, vancomycin MIC ≥ 32µg/mL, nitrofurantoin MIC ≥ 128µg/mL and high-level gentamicin resistance at MIC ≥ 500µg/mL.

**Table 2.**
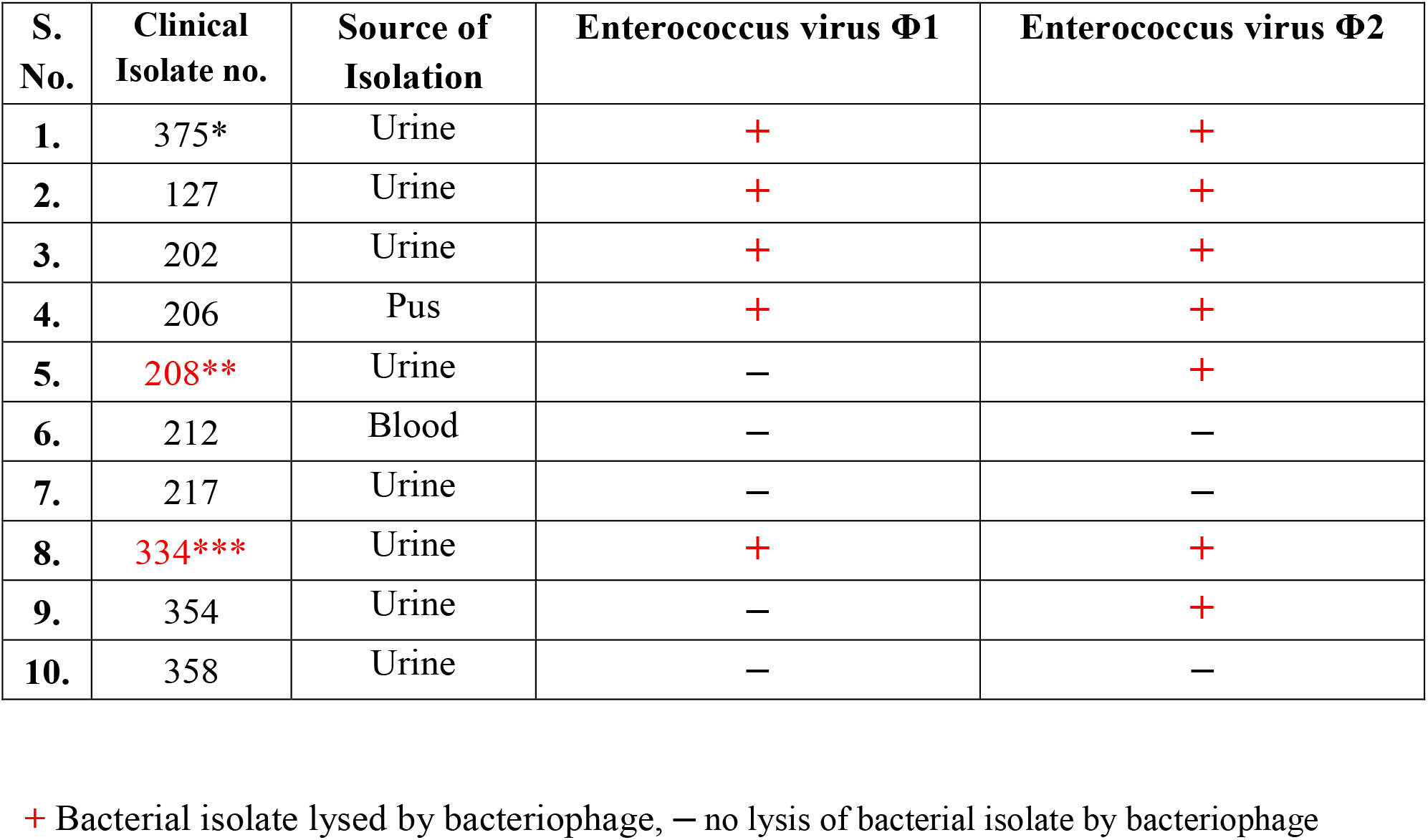
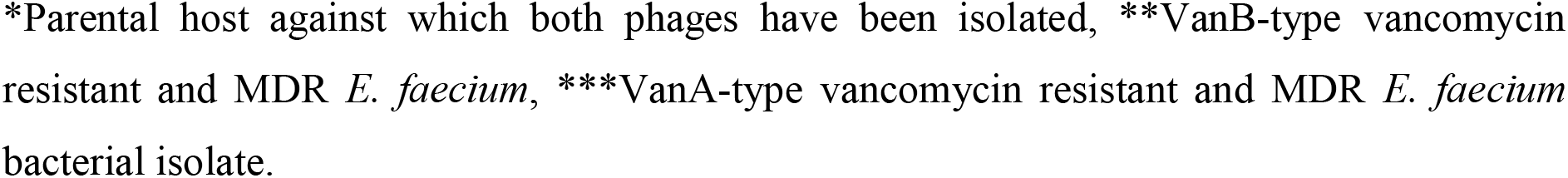
Host range analysis of Enterococcus virus Φ1 and Enterococcus virus Φ2 against MDR clinical isolates of *E. faecium*.

| S. No. | Clinical Isolate no. | Source of Isolation | Enterococcus virus Φ1 | Enterococcus virus Φ2 |
| --- | --- | --- | --- | --- |
| 1. | 375* | Urine | + | + |
| 2. | 127 | Urine | + | + |
| 3. | 202 | Urine | + | + |
| 4. | 206 | Pus | + | + |
| 5. | 208** | Urine | — | + |
| 6. | 212 | Blood | — | — |
| 7. | 217 | Urine | — | — |
| 8. | 334*** | Urine | + | + |
| 9. | 354 | Urine | — | + |
| 10. | 358 | Urine | — | — |

+ Bacterial isolate lysed by bacteriophage, — no lysis of bacterial isolate by bacteriophage
\*Parental host against which both phages have been isolated, \*\*VanB-type vancomycin resistant and MDR *E. faecium*, \*\*\*VanA-type vancomycin resistant and MDR *E. faecium* bacterial isolate.

**Fig. 3.**
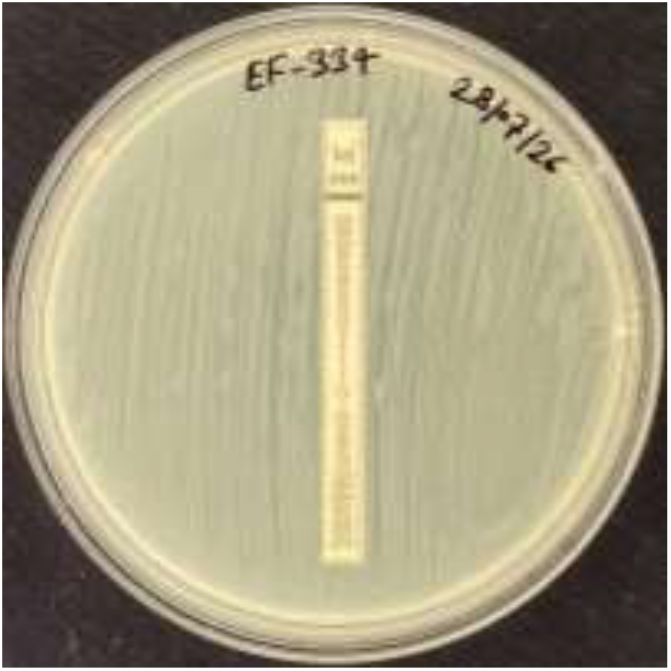
The *E. faecium* 334 isolate showing high vancomycin resistance (>256 µg/mL) by Ezy MIC™ strip.

**Fig. 4.**
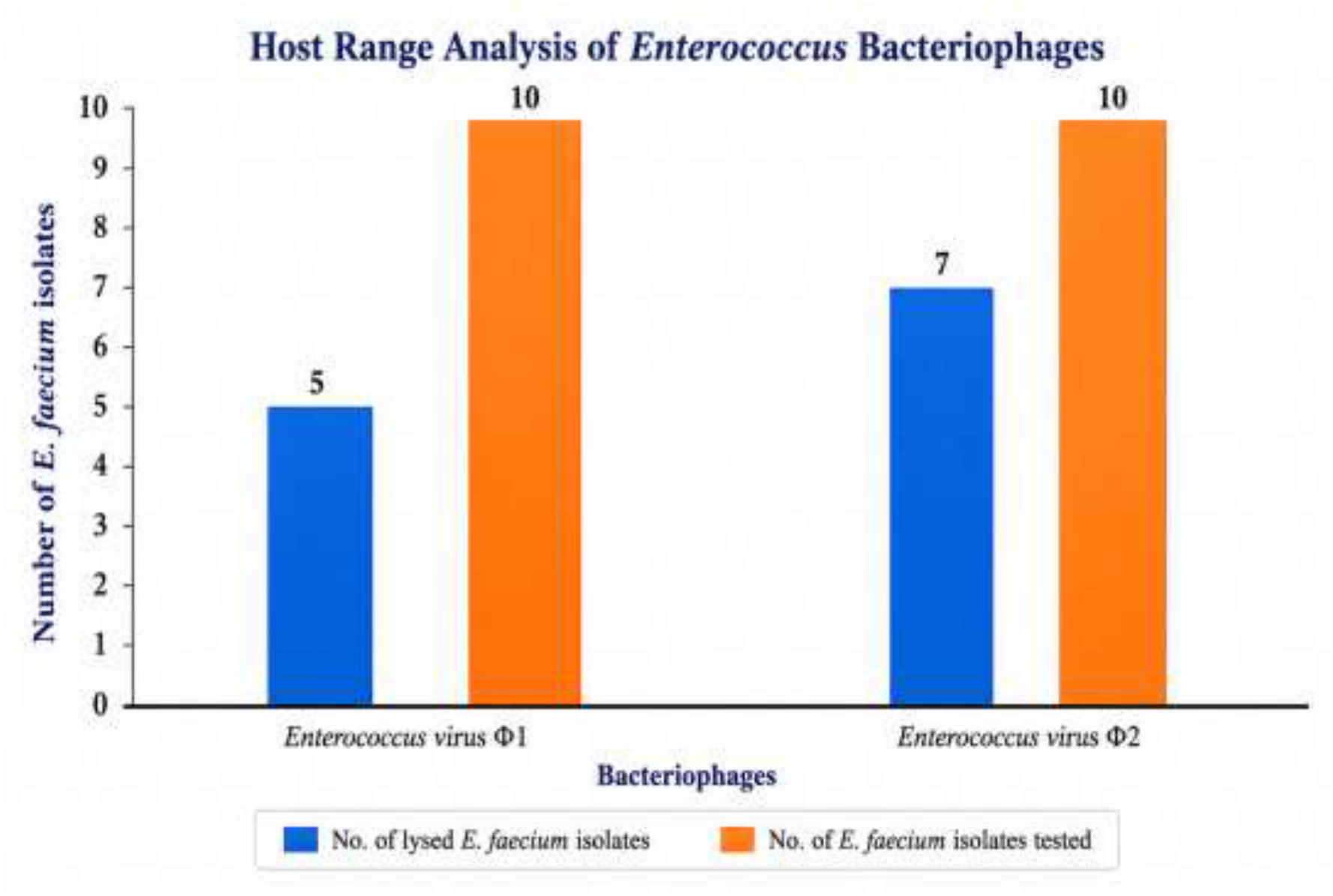
Host range analysis revealed that Enterococcus virus Φ1 exhibited lytic activity against 5 out of the 10 tested MDR *Enterococcus faecium* strains, whereas Enterococcus virus Φ2 demonstrated a comparatively broader host range, lysing 7 of the 10 isolates.

*E. faecium* isolate 334 showed resistance to both vancomycin and teicoplanin by the VITEK 2 system (Table 1). However, since VITEK 2 only reports result up to ≥32 µg/mL, the high-level vancomycin resistance of isolate 334 was confirmed using the Ezy MIC™ strip, which showed an MIC >256 µg/mL (Fig. 3). Based on the susceptibility patterns to vancomycin (Fig. 3) and teicoplanin (Table 1), *E. faecium* 334 was classified as the VanA phenotype, while *E. faecium* 208 was classified as the VanB phenotype, owing to resistance to vancomycin (MIC ≥32 µg/mL) combined with susceptibility to teicoplanin (MIC = 1 µg/mL).

### Phage Time-Kill Assay

The phage-time kill assay demonstrated a strong and sustained antibacterial effect of the bacteriophage against the host bacterium. Phage kill kinetics of Enterococcus virus Φ1 treatment against the parent MDR *E. faecium* strain EF375 over a 24 h period at 37 °C. The negative control (red; media only, no bacteria) maintained a stable, low OD throughout the 24 h period, confirming the absence of contamination and the stability of the assay conditions. The untreated bacterial control (blue) exhibited rapid exponential growth, followed by a typical stationary phase and a slight decline at later time points. In contrast, the phage-treated culture (green) showed only a modest initial increase in OD to approximately 0.25 during the first 2-3 h, after which the bacterial population declined and stabilized close to the negative control level for the remainder of the experiment, corresponding to an approximately 80-82% reduction in bacterial growth relative to the untreated control at 24 h (**Fig. 5. a)**.

**Fig. 5.**
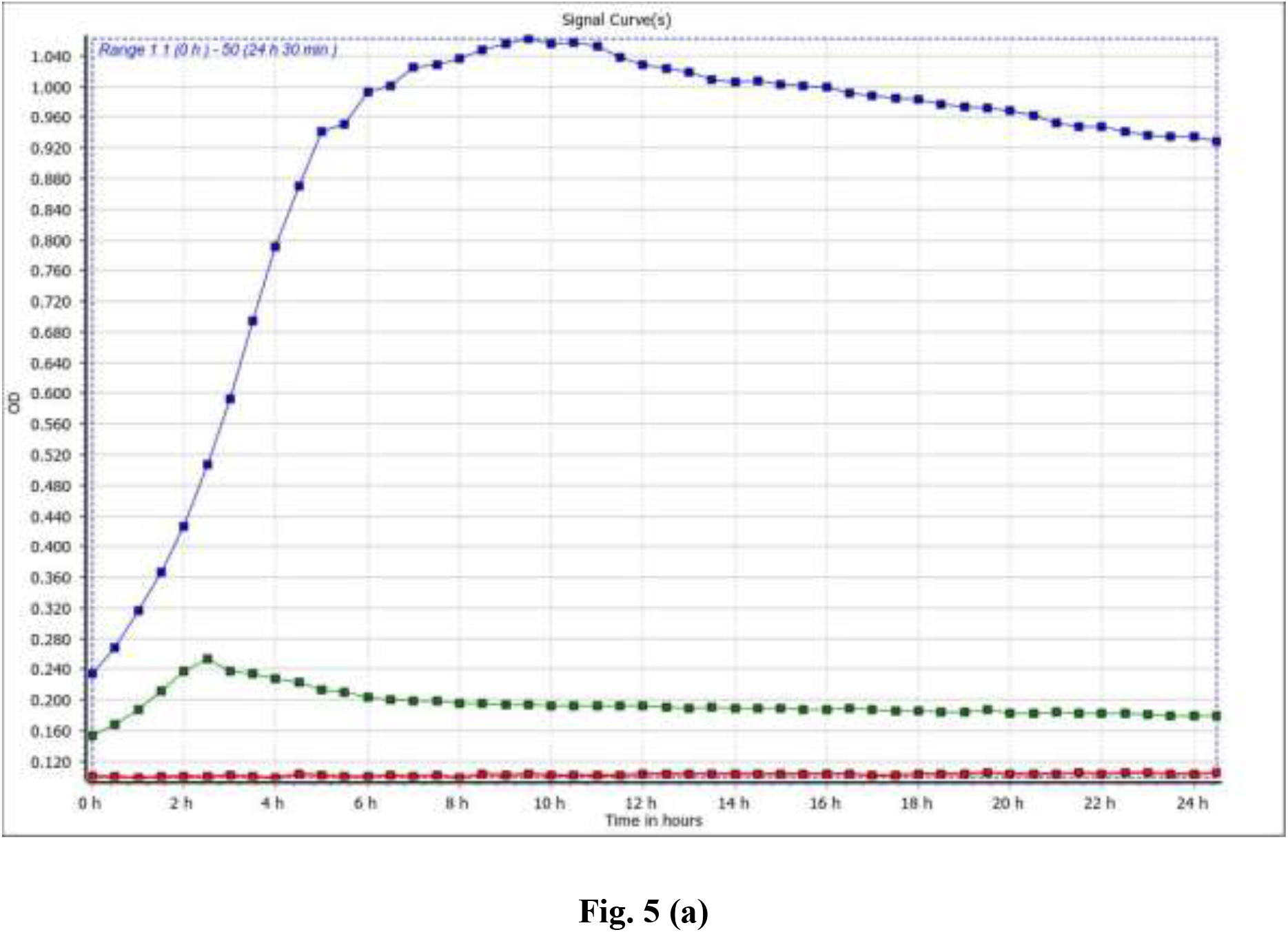
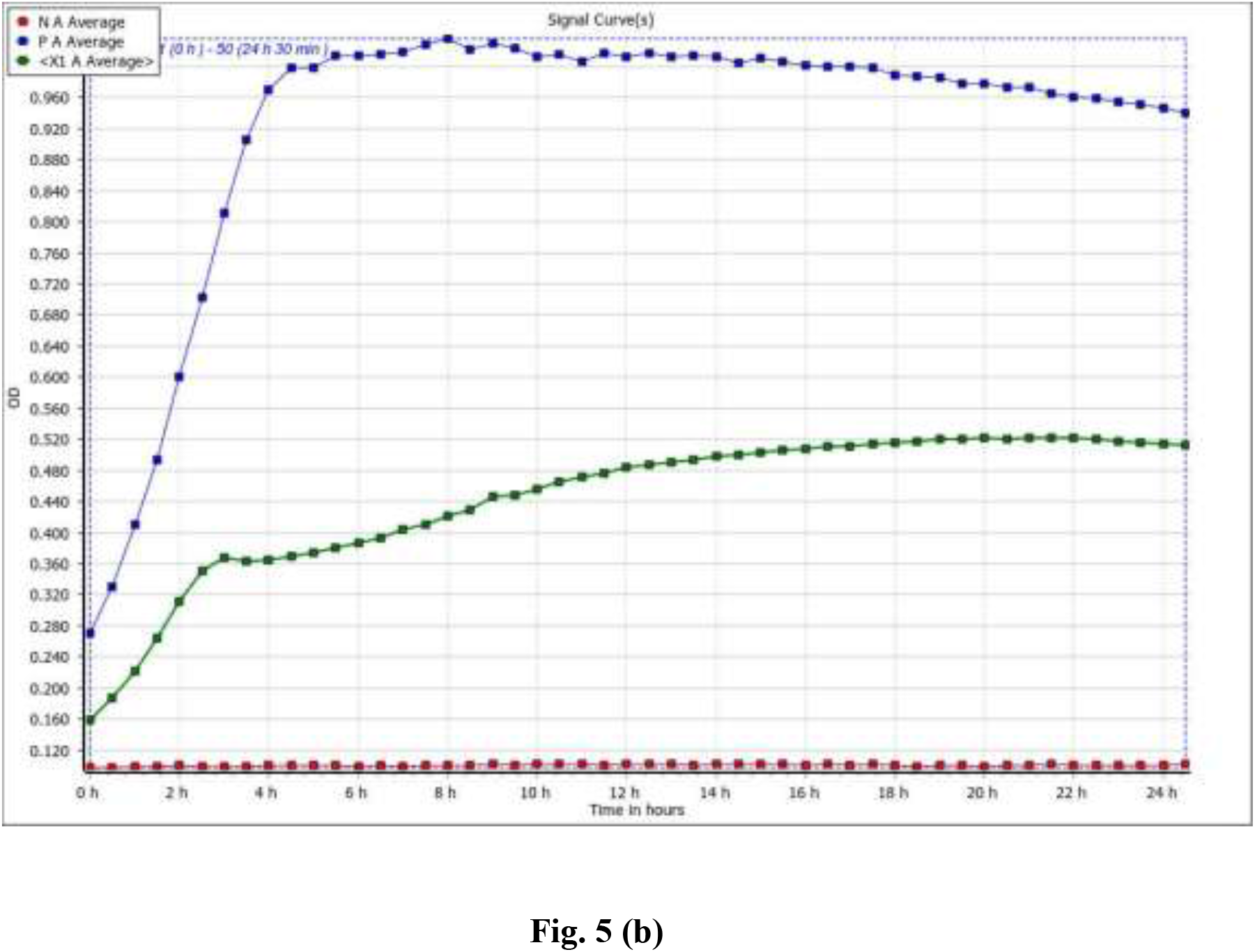
Phage kill kinetics with MDR parent host EF375 in liquid culture. The X-axis represent time interval represented in hours and Y-axis represent optical density (OD) at 600 nm and. The OD was measured at every 30 min up to 24 h. The untreated bacterial positive control (blue) exhibited rapid exponential growth, the red colour indicated the media only negative control. **(a**) In the presence of Enterococcus virus Φ1, the phage-treated culture (green) showed only a slight initial increase in bacterial density during the first 2-3 h, followed by suppression of bacterial growth. **(b)** In presence of Enterococcus virus Φ2, phage-treated culture (green) showed significant suppression of bacterial growth.

Another kill kinetics study with Enterococcus virus Φ2 with their parent MDR *E. faecium* strain EF375, the negative control (red; media only, no bacteria) again remained stable at a low OD throughout the assay, confirming the absence of contamination and the stability of the assay conditions. The untreated bacterial control (blue) again showed rapid exponential growth, whereas the phage-treated culture (green) showed markedly reduced growth throughout the experiment, with OD increasing only gradually to approximately 0.52 by 20-22 h, remaining substantially lower than the untreated control and corresponding to a nearly 48-50% reduction in bacterial growth relative to the untreated control at 24 h (**Fig. 5. b)**.

## Discussion

Vancomycin-resistant *E. faecium* is a major opportunistic pathogen that colonises in the gastrointestinal tract and subsequently can cause serious healthcare infections including urinary tract infections, endocarditis and bacteraemia and wound infections [44]. These pathogens show resistance to multiple classes of antibiotics and are frequently associated with persistent, chronic and recurrent infections [45]. The ability of *E. faecium* to persist in hospital environments, rapidly acquire multidrug resistance and ability to cause severe nosocomial infections has established it as a major healthcare-associated pathogen, underscoring the urgent need to develop novel preventive and therapeutic strategies against vancomycin-resistant *E. faecium* [46]. In recent years, bacteriophage therapy has emerged as a promising alternative for the treatment of infections caused by vancomycin-resistant *E. faecium* owing to its ability to specifically target antibiotic-resistant bacteria [47,48]. Although bacteriophages against MDR *E. faecium* have previously been reported from India and other countries [42, 47, 48], to the best of our knowledge, this is the first study describing bacteriophages with lytic activity against clinical VREfm isolates from India.

In the present study, two lytic *Enterococcus* virus NIMS_EF375_N69_P9 (Φ1) and NIMS_EF375_N74_P12 (Φ2) were isolated from sewage water samples against the MDR *E. faecium* (EF375) clinical isolate, cultured from urine sample. TEM analysis revealed that both phages have a regular icosahedral head and a long non-contractile tail, characteristic of tailed bacteriophages belonging to the class *Caudoviricetes*, with a *Siphovirus*-like morphology. These findings are similar to prior reports, mentioning the *Siphovirus* morphotype for most of the *E. faecium* phages [49-50]. Both phages showed a broad host range against MDR clinical isolates of *E. faecium*. Specifically, Φ2 presented a comparatively broader host range, lysing 7 out of 10 clinical strains (including the parental propagation host; 6 of 9 non-parental isolates). In evaluating phage activity against VREfm strains, we observed that Phage Φ2 was able to lyse both VanA-type VREfm (EF 334) and VanB-type VREfm (EF 208), while Phage Φ1 showed activity against only VanA-type VREfm (EF 334) strain. Most of the previously reported *E. faecium* phages have shown relatively narrow host ranges, although a few broad-host-range phages have been reported [49,50,51]. A broad host range is a desirable characteristic for therapeutic phages. Although the number of isolates tested in the present study was smaller than in previous reports, the broad lytic spectrum observed, particularly for Φ2, highlights its potential as a promising candidate for the treatment of VREfm *E. faecium* infections, particularly given its activity against both VanA- and VanB-type VREfm isolates.

In the phage time-kill assay, both phages showed a significant reduction in bacterial culture in the presence of bacteriophage; however, their behavior differed. The Φ1-treated culture inhibited bacterial growth by up to 80%, while Φ2 reduced bacterial growth by approximately 50%, compared to the phage-free culture. The lower lytic efficiency of Φ2 may be attributed to a longer latent period, lower burst size, or the emergence of phage-resistant bacterial cells. Interestingly, the host range and lytic pattern in the time-kill assay showed an inverse relationship: Φ1 exhibited higher lytic efficiency but a narrower host range, while Φ2 produced less efficient lysis but a broader host range. Such trade-offs between host-range breadth and lytic potency have also been reported for other phage-host interactions, and may be relevant when selecting candidates for phage-cocktail formulations to achievecomplementary strengths [52,53]. Viable bacterial counts (CFU/mL) alongside extended monitoring beyond 24 h would help clarify these dynamics and confirm whether the Φ2-treated population reflects residual susceptible cells or true phage-resistant mutants. Taken together, these findings support the therapeutic potential of both bacteriophages for controlling VREfm and MDR *E. faecium* infections.

Phage therapy against *E. faecium* holds considerable promise, particularly in light of the increasing prevalence of VREfm and MDR strains. However, several challenges must be addressed before its widespread clinical implementation. These include the phage cocktails with broaden host coverage, the emergence of phage-resistant bacterial population and the complexities associated with large-scale production, standardization and regulatory approval of phage-based therapeutics. Moreover, the therapeutic efficacy of phage treatment may be enhanced through prolonged treatment regimens and use of phage cocktails and synergistic combinations with conventional antibiotics [48, 54]. The present study has certain limitations. The host range was evaluated against a relatively small number of clinical isolates, and whole-genome sequencing and in vivo therapeutic efficacy studies were not performed. Further investigations involving larger isolate collections and animal infection models are required to validate the therapeutic potential of these bacteriophages.

## Acknowledgements and Funding

The authors are thankful to Indian Council of Medical Research (ICMR), New Delhi, India to provide funds to establish Centre for Advance Research for Bacteriophage Research and Therapy (Grant No. ICMRCARREP-2023-000259).

## AI Disclosure Statement

Claude (Anthropic) was used exclusively for language editing, formatting of the reference list and improving the readability of this manuscript. The authors are fully responsible for the content presented in the manuscript.

